# Single cell transcriptomic analysis of bloodstream form *Trypanosoma brucei* reconstructs cell cycle progression and differentiation via quorum sensing

**DOI:** 10.1101/2020.12.11.420976

**Authors:** Emma M. Briggs, Richard McCulloch, Keith R. Matthews, Thomas D. Otto

**Affiliations:** Institute for Immunology and Infection Research, School of Biological Sciences, University of Edinburgh, Edinburgh, UK; Wellcome Centre for Integrative Parasitology, Institute of Infection, Immunity and Inflammation, University of Glasgow, Glasgow, UK

## Abstract

The life cycles of African trypanosomes are dependent on several differentiation steps, where parasites transition between replicative and non-replicative forms specialised for infectivity and survival in mammal and tsetse fly hosts. Here, we use single cell transcriptomics (scRNA-seq) to dissect the asynchronous differentiation of replicative slender to transmissible stumpy bloodstream form *Trypanosoma brucei*. Using oligopeptide-induced differentiation, we accurately modelled stumpy development *in vitro* and captured the transcriptomes of 9,344 slender and stumpy stage parasites, as well as parasites transitioning between these extremes. Using this framework, we detail the relative order of biological events during development, profile dynamic gene expression patterns and identify putative novel regulators. Using marker genes to deduce the cell cycle phase of each parasite, we additionally map the cell cycle of proliferating parasites and position stumpy cell cycle exit at early G1, with subsequent progression to a distinct G0 state. We also explored the role of one gene, ZC3H20, with transient elevated expression at the key slender to stumpy transition point. By scRNA-seq analysis of ZC3H20 null parasites exposed to oligopeptides and mapping the resulting transcriptome to our atlas of differentiation, we identified the point of action for this key regulator. Using a developmental transition relevant for both virulence in the mammalian host and disease transmission, our data provide a paradigm for the temporal mapping of differentiation events and regulators in the trypanosome life cycle.

## Introduction

African trypanosome parasites cause both human^1^ and animal^2^ trypanosomiases and are transmitted between hosts across sub-Saharan Africa by tsetse flies. During its life cycle *Trypanosoma brucei* undergoes several developmental transitions, comprising changes in nutrient-specific metabolism, morphology, organelle organisation and structure, and stage-specific surface protein expression^3^, facilitating parasite survival and transmission. In the mammalian host, long slender bloodstream forms replicate extracellularly, increasing in numbers to trigger differentiation into short stumpy bloodstream form parasites via a quorum sensing (QS) process^4,5^, with ill-defined intermediate forms between these morphological extremes^6,7^. Stumpy forms remain arrested in the cell cycle^8^ until ingested by a feeding tsetse fly, where they are pre-adapted to survive in the midgut^9,10^. Here, stumpy forms undergo a further differentiation event and re-enter the cell cycle as tsetse-midgut procyclic forms^9,11^.

Slender and stumpy forms differ at both the transcript^12–17^ and protein level^18,19^, as do stumpy and procyclic parasites^15–17,19^. Reflecting their metabolism, slender forms show high levels of transcripts encoding glycosomal components (specialist organelles housing glycolytic enzymes)^9^, whereas stumpy parasites upregulate transcripts related to a maturing mitochondrion as they prepare for the tsetse midgut. This allows for metabolism of pyruvate, as well as proline and threonine, to generate ATP in low glucose conditions^9,13–15,20^. Consistent with exit from the cell cycle, stumpy parasites down-regulate histone, DNA replication/repair, translation and cytoskeleton-related transcripts ^15^. Additionally, PAD (Proteins Associated with Differentiation) transcripts are upregulated in stumpy forms and are required for further development into procyclics^21^. Transcripts encoding EP and GPEET repeat procyclin surface proteins expressed in tsetse midgut forms are also elevated in stumpy forms, whereas variant surface glycoprotein (VSGs) transcripts, required for immune evasion by the parasite in the mammal, are reduced. Transcript analysis of *T. brucei* parasites isolated during parasitaemia *in vivo* suggested some of these changes occur in early differentiating parasites, before morphologically detectable stumpy forms dominate at the peak of parasitemia^22^.

QS based development between slender and stumpy forms has been recently characterised, identifying several factors involved in detecting the differentiation stimulus^23^, signal propagation^24,25^ and implementation of cellular changes^24,26–28^. Yet, understanding the detailed developmental progression toward stumpy cells has been hampered by the asynchrony of this differentiation step, as has the relationship of differentiation regulatory genes to the various biological events involved. However, single-cell RNA sequencing (scRNA-seq) offers the opportunity to study individual cells in a heterogenous population, to identify rare cell types and decipher complex and transient developmental processes^29–31^. Recently, scRNA-seq has been used to study antigenic variation in *T. brucei*^32^, as well as to describe the diversity of parasites in the tsetse fly salivary gland^33^. The latter study revealed early and late stages of metacyclic development, previously indistinguishable by population-based RNA-seq^34^, highlighting the differing expression of surface proteins within the developing population^33^.

Here, we applied scRNA-seq to analyse 9,344 differentiating parasites progressing from bloodstream slender, through intermediate, to stumpy parasites *in vitro* using oligopeptide-rich bovine brain heart infusion (BHI) broth^23^, deriving a temporal map of this transition at the transcript level based on individual cells. Detailed analysis of the associated expression patterns revealed the absence of a discrete “intermediate” transcriptome, mapped the relative timing of biological events during differentiation, including exit from the cell cycle specifically prior to late G1, and identified novel genes regulated during the transition. Moreover, scRNA-seq analysis of a null mutant for one important regulator elevated at the slender to stumpy transition, ZC3H20^26,27^, precisely mapped where development fails in its absence in molecular terms. In combination, this provides a paradigm for the temporal mapping of developmental events and regulators during the parasite’s dynamic differentiation programme in its mammalian host.

## Results

### scRNA-seq identifies transcriptionally distinct long slender and short stumpy form *T. brucei*

To model stumpy development *in vitro*, pleomorphic *T. brucei* EATRO 1125 AnTa1.1 90:13 slender parasites were treated with oligopeptide-rich BHI broth, able to induce *T. brucei* bloodstream form differentiation in a titratable manner^23^. In the presence of 10% BHI, parasites underwent growth arrest (Fig. S1a), increased expression of the stumpy marker protein PAD1^35^ (Fig. S1b), and increased the percentage of parasites containing one copy of the nucleus and one copy of the kinetoplast network (1N1K), indicating cell cycle accumulation in G1/G0 and differentiation into stumpy forms^8^ (Fig. S1c). After 72 hr, 72.5% of cells expressed PAD1 (Fig. S1b) and 89.3% were in the 1N1K cell cycle configuration (Fig S1c). To capture the transcriptomes of slender, intermediate and stumpy *T. brucei*, we combined parasites after 0, 24, 48 or 72 hr of 10% BHI treatment in equal numbers. 15,000 cells of this heterogenous pool were then subjected to scRNA-seq using the Chromium Single Cell 3’ workflow (10X Genomics) and Illumina sequencing^36^. Two independent biological replicates (WT 1 and WT 2) were generated and, after filtering to remove transcriptomes of poor quality or likely doublets, 9,344 cells remained (5,793 and 3,551, respectively; Fig. S2 and Supplementary data 1). The transcripts of 8,757 genes were captured in at least 5 cells in both replicate experiments (10 cells total), with medians of 1,051 and 1,439 genes detected per cell, respectively. Cells from the two replicate experiments were integrated and UMAP (Uniform Manifold Approximation and Projection^37^) was used to visualise the relationship between individual *T. brucei* transcriptomes in low dimensional space, where variation between transcriptomes dictates the space between cells (Fig. 1a, b, d). Cells from WT 1 and WT 2 experiments overlapped (Fig. 1a), indicating the capture of reproducible cell types in each replicate. Clustering analysis identified four distinct groups containing transcriptionally similar cells (Fig. 1b; Supplementary data 2), each appearing in comparable proportions per replicate (Fig. 1c).

**Figure 1.**
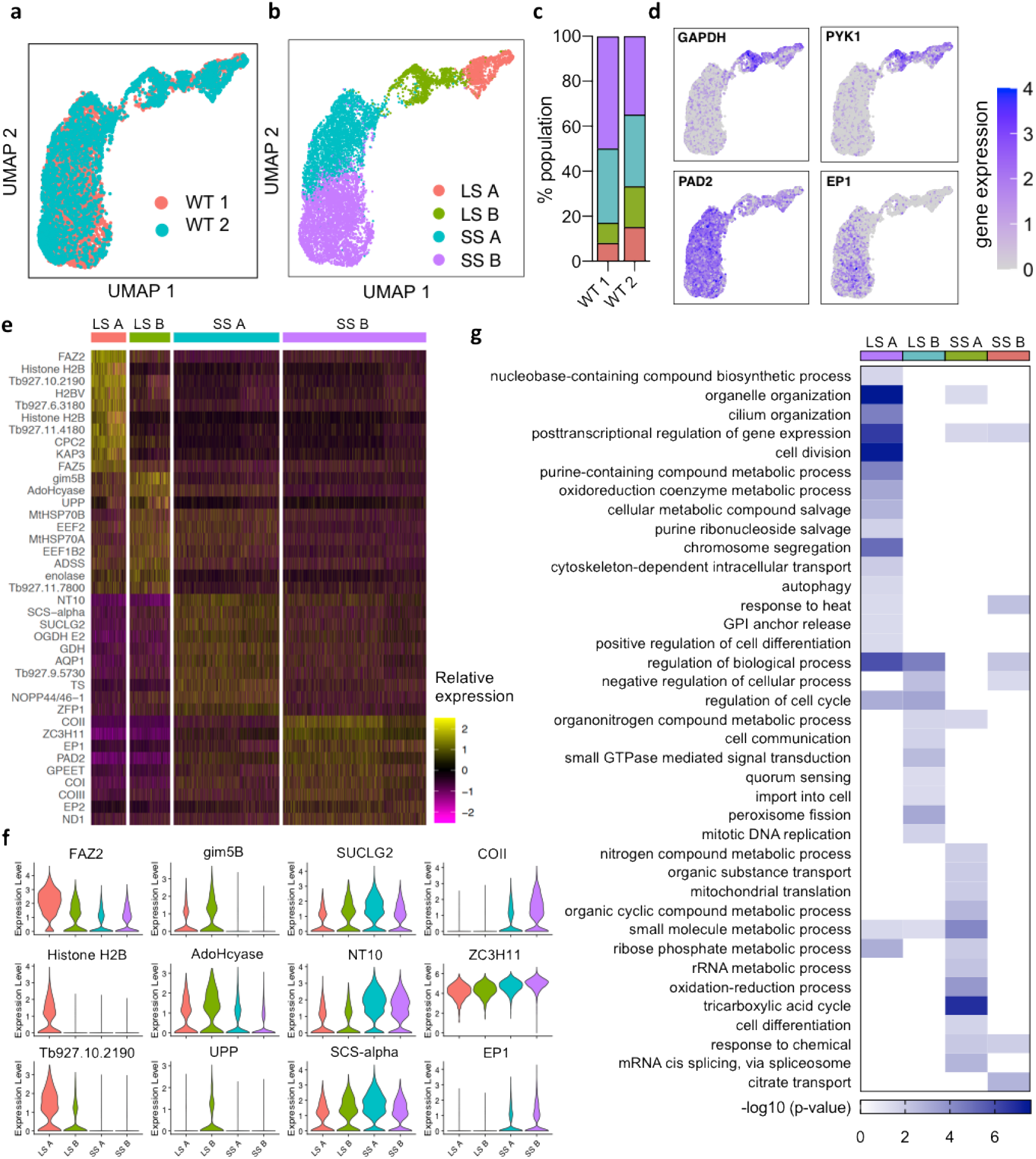
Sequencing of individual *T. brucei* transcriptomes during bloodstream differentiation *in vitro*. **a)**. Low dimensional plot (UMAP) of each cell after filtering. Each point is the transcriptome of one cell positioned according to similarity with neighboring transcriptomes, coloured by replicate experiment. **b)** UMAP of WT parasites from both replicates, coloured by cluster: Long slender (LS) A, LS B, short stumpy (SS) A and SS B. **c)** Percentage of parasites in each cluster for each replicate experiment. **d)** UMAP of integrated WT parasites coloured by transcript counts for two SL marker genes (GAPDH; Tb927.6.4280 and PYK1; Tb927.10.14140) and stumpy marker genes (PAD2; Tb927.7.5940 and EP1; Tb927.10.10260). Scale shows raw transcript count per cell. **e**) Heatmap showing relative expression of the top 10 unique maker genes of each cluster identified in **c**. Each row is one gene coloured by relative expression. Where no gene name or symbol was available, the gene ID is shown. Each column is one cell grouped according to cluster. **f)** Violin plots showing the expression of each of the top 3 unique marker genes per cell, divided by cluster. X-axis shows the raw transcript count per cell. **g**) Gene ontology (GO) enrichment for biological process linked with maker genes for each cluster. Scale shows the −log(adjusted p-value) for each term enrichment per cluster.

Slender- and stumpy-like cells were clearly identifiable by the expression of marker genes: slender-associated glycolytic genes GAPDH and PYK1^38^; and the stumpy markers, PAD2^21^ and EP1 procyclin^13^ (Fig. 1d). Differential expression analysis of transcripts between the slender A, slender B, stumpy A and stumpy B clusters identified 519 marker genes (adjusted p-value < 0.05, logFC > 0.25); relative expression of the top unique markers is plotted in Fig. 1e. Markers of slender A and slender B overlapped extensively (Supplementary data 2); 178 slender marker genes were upregulated in both slender A and slender B relative to stumpy clusters, including glycolytic genes (for example, hexokinase, phosphoglycerate kinase, glucose-6-phosphate isomerase), genes involved in cytoskeleton organisation (e.g. beta tubulin, cytoplasmic dynein 2 heavy chains 1 and 2), cell cycle regulating genes (e.g. cyclin 10, cyclin-like F-box protein 2 and T-complex protein 1 subunits gamma and delta) and RNA-binding protein 10 (RBP10), which is a positive regulator of bloodstream form transcripts^39,40^. Markers unique to slender A or slender B (183 and 95, respectively) were generally related to the cell cycle phase of these cells and are discussed in detail below. The top differentially expressed markers of slender A were FAZ2 (flagellum attachment zone protein 2), CPC2 (chromosomal passenger complex 2) and histone H2B (Fig 1e and f). Glycosomal membrane protein gim5B, a putative S-adenosylhomocysteine hydrolase (AdoHcyase) and uridine phosphorylase (UPP) were the top markers of slender B (Fig. 1e and f). 55 genes were upregulated in stumpy A cells relative to the other clusters, including purine nucleoside transporter NT10 (known to be associated with stumpy forms^41,42^), succinyl-CoA synthetase alpha subunit (SCS-alpha), and succinyl-CoA ligase [GDP-forming] beta-chain (SUCLG2) (Fig. 1e and f). Just 9 genes significantly distinguished stumpy B cells, including four encoded by the mitochondrial genome: cytochrome oxidase subunits I-III (COI, COII, COIII) and NADH dehydrogenase subunit 1 (ND1) (Fig. 1e,f).

Gene ontology (GO) term enrichment analysis revealed the association of each cluster’s marker genes with distinct biological processes (Fig. 1g). Several terms relating to cell cycle processes were enriched in slender A marker genes: organelle and cilium organisation, chromosome segregation, and cell division. Cell cycle regulation genes, including Cytokinesis Initiation Iactor 1 (CIF1)^43^ and 14-3-3 protein 1 (14-3-3-I)^44^, glycosylphosphatidylinositol-specific phospholipase C (GPI-PLC), involved with GPI anchor release^45^, and a positive regulator of differentiation, ZC3H20^26,27,46^, were all upregulated in slender A and slender B cells compared to stumpy cells. Slender B was also associated with the genes implicated in quorum sensing, including protein phosphatase 2C (PP2C) and Trichohyalin^24^, as well as genes putatively involved in cell communication (Ras-related protein Rab5A^47^ and thimet oligopeptidase). Stumpy A marker genes were associated with the TCA cycle (mitochondrial malate dehydrogenase, two 2-oxoglutarate dehydrogenase E1 component encoding genes, and succinyl-CoA ligase [GDP-forming] beta-chain^48^), oxidation-reduction process (dihydrolipoyl dehydrogenase^4,49–51^ and glutamate dehydrogenase^52^), rRNA metabolism (splicing factor TSR1^53^, nucleolar RNA-binding protein NOPP44/46-1^54–56^ and Lupus LA protein homolog^57^), and cell differentiation (zinc finger protein 2; ZFP2^58^). GO term analysis of stumpy B marker genes was limited due to their small number but included post-transcriptional regulators of gene expression, due to the presence of ZC3H11, which is also involved in heat shock response^59^, and PAD2, which is involved in detecting the differentiation to procyclic forms stimulus via citrate transport^21^.

Taken together, the above clustering analysis revealed distinct slender and stumpy clusters, with significant variation within each population. Interestingly, a distinct cluster representative of a discrete “intermediate” stage transcriptome between slender and stumpy forms was not evident.

### Trajectory analysis of long slender to short stumpy differentiation

As clustering analysis highlighted the transition from slender and stumpy cells involved overlapping gene expression and GO term association, we conducted trajectory inference and pseudotime analysis to study gene expression changes during stumpy development in detail. Individual cells were replotted as a PHATE (Potential of Heat-diffusion for Affinity-based Transition Embedding) map (Fig. 2a-c), which captures the local and global structure of high-dimensional data to preserve the continual progression of developmental processes^60^. Here, slender A and slender B clusters remained clearly separate, whereas stumpy A and stumpy B showed more extensive overlap (Fig. 2a). A linear trajectory starting from slender A cells was identified (Fig. 2b). Slender and stumpy marker gene (GAPDH, PYK1, PAD2 and EP1) expression across the trajectory confirmed capture of the transition from slender to stumpy forms (Fig. 2c). 2001 genes were identified as differentially expressed as a function of pseudotime (p value < 0.05, fold change > 2) and were grouped into 9 modules (A-I) of co-expressed genes that showed similar patterns of expression across differentiation (Fig. 2d; Supplementary data 2). Of these, 1,335 genes were previously found to be significantly (adjusted p-value < 0.05) differentially expressed between slender and stumpy enriched populations of *T. brucei* isolated from low and peak parasitaemia *in vivo*, respectively^12^, confirming the physiological relevance of *in vitro*, oligopeptide induced, differentiation (Fig. 2e). Proportionally fewer genes in modules A (transiently down regulated) and F (transiently upregulated) had been identified in bulk RNA-seq data as differentially expressed (33.3% and 49%, respectively) compared to the remaining modules (59.1 - 78.9%), highlighting the ability of single cell analyses to reveal transient events in an asynchronous developmental trajectory.

**Figure 2.**
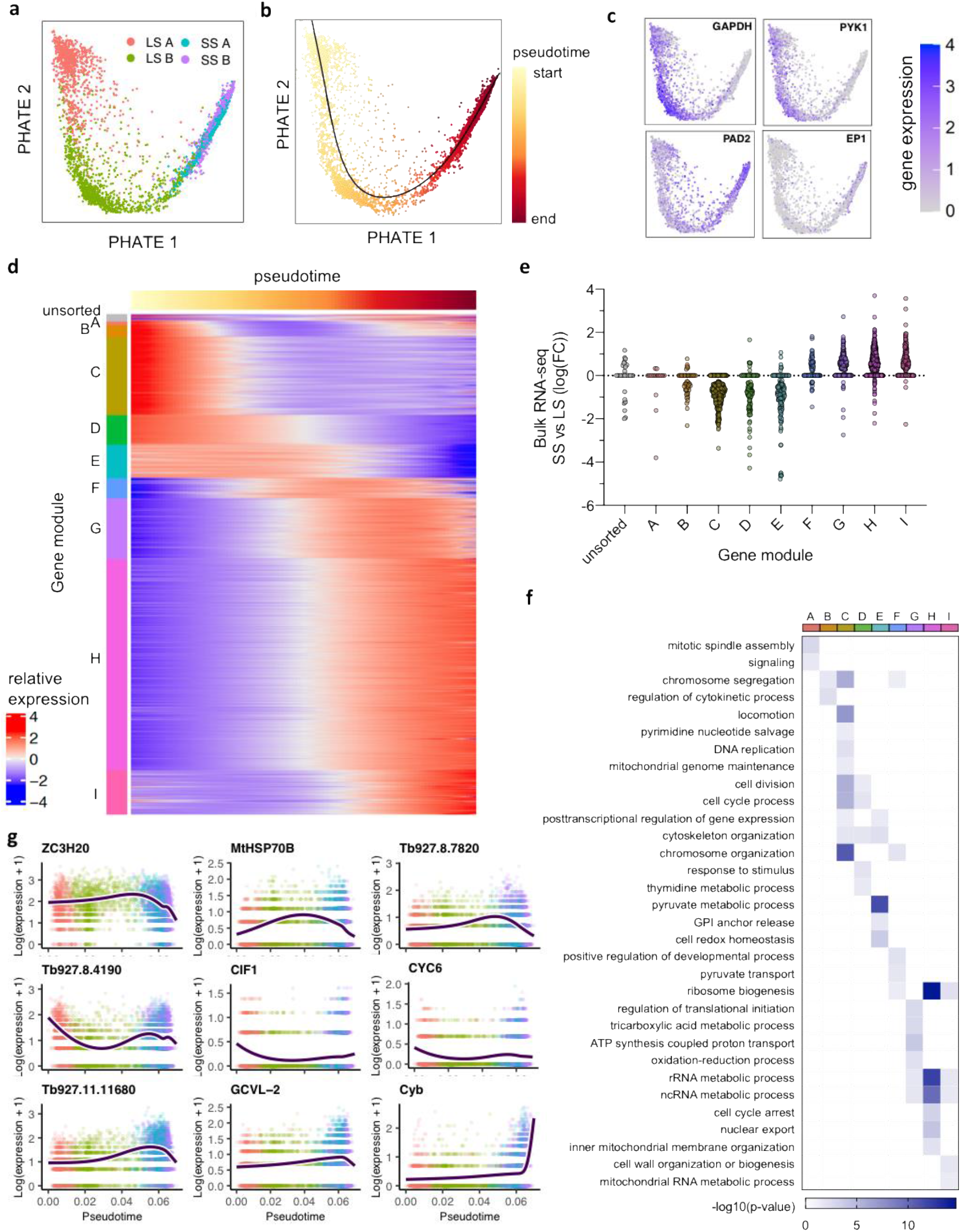
Pseudotime analysis reveals dynamic gene expression in slender to stumpy differentiation. PHATE plots of individual parasite transcriptomes coloured by (**a**) cluster identity, (**b**) pseudotime values and (**c**) raw marker gene transcript count as in **1e**. **d**) Heatmap plotting relative expression of genes significantly (>0.05 adjusted p value, FC > 2) associated with the trajectory (2001 genes). Top track shows pseudotime. Genes are clustered by expression pattern over pseudotime into 9 modules of co-expressed gene, indicated to the left. Unsorted genes are indicated (grey). **e**) Fold change of differentiation associated genes in bulk in vivo-derived RNA-seq data, comparing stumpy (peak parasitemia) and slender (low parasitemia). Each point is one gene, grouped and coloured according to the co-expressed module identified in d. **f**) Biological progress gene ontology (GO) term analysis of differentiation associated genes grouped by co-expressed module. **g**) Gene expression (log(transcript count +1)) across pseudotime from slender to stumpy differentiation of 9 genes identified as transiently upregulated (ZC3H20, MtHSP70B, Tb927.8.7820), transiently down regulated (Tb927.8.4190, CIF1, CYC6) or unsorted (Tb927.11.11680, GCVL-2, Cyb). Each point is one cell coloured by cluster as in **a**. Dark blue line is smoothed average expression across pseudotime.

GO term enrichment for biological processes associated with each gene module revealed the relative order of biological events during slender to stumpy development (Fig. 2f). Processes upregulated at the start of the trajectory included chromosome segregation and regulation of cytokinesis (module B), and these were then followed by cell division (modules C and D), indicating the progression of the later stages of the cell cycle. Module D also included genes linked to stimulus response, including PKA-R (previously identified as a potential signal transducer during stumpy development in response to hydrolysable-cAMP^24^), cAMP-specific phosphodiesterases 1 and 2 (PDEB1 and PDEB2), the known differentiation regulator RBP7B^24,25^, DHFR-TS, an enzyme required for thymine synthesis^61^, and mitotic cyclin CYC8^62^. Module E genes, which are broadly expressed across the slender parasites and peaked in expression slightly later in pseudotime, include GPI-PLC, which releases the VSG coat via hydrolysis of the GPI-anchor^45^, and genes associated with cytoskeleton organisation (beta tubulin, actin A, cytoskeleton associated proteins CAP51V and CAP5.5V^63,64^). Module F consisted of transiently upregulated genes, including the known stumpy and procyclic developmental regulator ZC3H20^26,27,46^, and MCP1, a pyruvate transporter present in the mitochondrial membrane^65^. Module G-I genes peaked in the later stages of development and include further genes identified in a reverse-genetic screen for stumpy development factors: the chromatin regulator ISWI, a phosphoglycerate mutase protein, KRIPP14, APPBP1, and hypothetical proteins Tb927.11.300 and Tb927.11.1640^24^. Module G included genes encoding components of the TCA cycle: mitochondrial chaperone BCS1, and four ATP synthase genes (ATPF1A, ATPB and mitochondrial ATP synthase delta chain^48^). Protein coding genes associated with rRNA metabolism (n = 31) and ribosome biogenesis (n = 44) were also upregulated in the later stages of development, including 20 large ribosomal subunit components and 2 rRNA methyltransferases. These might be related to the translational preparedness exhibited by quiescent stumpy forms for development to procyclic forms and the resumption of translation^22^. Six genes linked to cell cycle arrest were identified in module H, all encoding copies of retrotransposon hotspot protein RHS4^66^. kDNA-encoded genes RPS12, ND1, COI-III, and NDH4^67^, were all increased in the later stages of development (modules G-I). Cytochrome B (Cyb), which was not sorted into a gene module, peaked at the very end of development (Fig. 2g). Beyond these annotated genes, 635 hypothetical genes were identified as differentially expressed during slender to stumpy differentiation, including predicted gene regulators^68^ Tb927.8.7820 (transiently upregulated), Tb927.8.4190 (transiently down regulated), and Tb927.11.11680, which peaks in stumpy cells (Fig. 2g).

Pseudotime analysis was able to identify novel genes differentially expressed during bloodstream form differentiation, as well as each gene’s detailed expression pattern. The relative timing of events, from proliferation (chromosome segregation, cytokinesis), cell cycle exit, cell remodelling, through to a maturing mitochondrion and expression of procyclin surface protein transcripts, can be inferred from these expression patterns. Additionally, the expression peaks of known and putative developmental regulators were identified relative to this progression.

### Transcript abundance during the bloodstream slender cell cycle

As replicating slender bloodstream form cells were captured in these experiments, we next asked if the scRNA-seq data could reveal greater detail than currently available on gene expression changes during the cell cycle. We first assigned each cell to a cell cycle phase using marker genes previously identified by bulk RNA-seq analysis^69^, (Fig. 3a, S3). Slender A and slender B cells were clearly grouped closer to cells of the same phase, with the parasites most distal to the stumpy A and stumpy B cells labelled as late G1, followed by S and G2/M phase cells. Slender B cells most proximal to stumpy A contained all four cell cycle phases, although early G1 cells were enriched here. Stumpy A and stumpy B cells were marked as a variety of all the cell cycle phases, indicating the phase of stumpy parasites cannot be clearly identified with these markers and stumpy parasites do not clearly reside with the G1 phase of actively replicating parasites.

**Figure 3.**
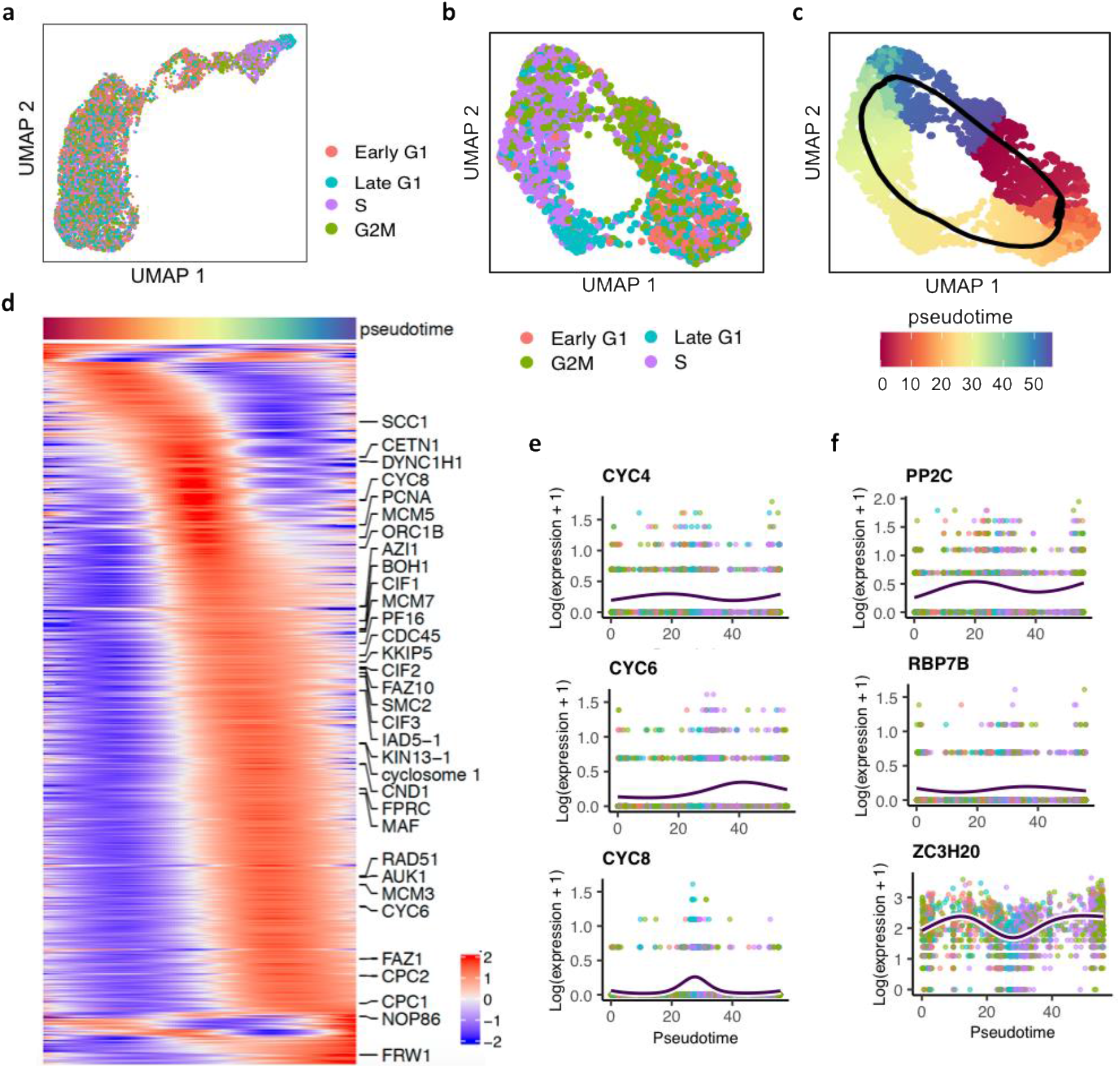
Identification of genes differentially expressed during the slender form cell cycle. **a)** UMAP of WT cells coloured by assigned cell cycle phase. **b)** UMAP of re-plotted slender A and slender B clusters of cells, using genes variable within the slender population. **c)** UMAP of slender cells coloured by assigned pseudotime value. The black line indicates the inferred circular trajectory. **d)** Heatmap of relative expression of genes significantly differentially expressed (p-value <0.05), with FC >2 over the cell cycle. Genes associated with the GO term “cell cycle process” are labelled. **e)** Expression (y-axis; log(expression +1)) of 3 cyclin genes (CYC4, CYC6 and CYC8) and cell cycle pseudotime (x-axis). Each point is one cell coloured by cell cycle phase and dark blue line shows average expression over pseudotime. **f)** Expression of three genes previously shown to regulate stumpy formation. Labels as in **e**.

Slender A and slender B clusters were next isolated and low dimensional projection of the data was repeated, resulting in these cells organising into a circular profile (Fig. 3b). Cells were clearly arranged by their cell cycle status within this ring, with late G1, S and G2/M phase cells distinguishable. Although early G1 cells were enriched between G2/M and late G1 cells, as expected, they overlapped with both neighbouring phases, indicating this stage was less well defined by the scoring. To assess gene expression changes during the cell cycle, we fitted a cyclical trajectory to this plot and assigned pseudotime values (Fig. 3c). Testing each gene for expression patterns associated with pseudotime identified 1,897 that were significantly (adj. p-value < 0.05) differentially expressed (Supplementary data 3). 854 of these changed in abundance by more than 2-fold (Fig. 3d). GO term enrichment revealed expected GO terms associated with the cell cycle; 33 genes associated with the term “cell cycle process” are highlighted in Fig. 3d. Amongst these 33 genes were several where protein levels or distribution have been shown to match the scRNA-seq predicted cell cycle timing of expression, including ORC1B^70^, AUK1^71^, PCNA^72^ and KKIP5^73^. Many further genes displayed cell cycle regulated expression that has not yet been explored (Supplementary data 3). For instance, two cyclin genes were identified, each with a distinct expression profile; CYC4 transcripts were more abundant before the appearance of late G1 and S phase cells and CYC6 peaked post-late G1. CYC8 peaked specifically during late G1, as previously documented by bulk RNA-seq analysis of cell cycle sorted populations^69^ (Fig. 3e).

In addition to genes driving the cell cycle, we identified three genes previously shown to be involved in stumpy development with differential expression patterns in slender cells (Fig. 3f). RBP7B^24^ expression increased in late G1 cells and persisted though to G2/M. PPC2, in contrast, showed the opposite pattern of expression, deceasing in late G1/S phase parasites. ZC3H20^26,27,46^ is highly abundant at the transcript level, yet dropped in expression in late G1/S phase *T. brucei*.

### ZC3H20 null parasites fail to differentiate in response to BHI

The above analysis highlighted differential expression patterns of several stumpy development regulators, including ZC3H20, which peaks in expression at the slender B to stumpy transition in pseudotime (Fig. 2g). As ZC3H20 has been previously shown to be required for differentiation *in vivo* and *in vitro* at high density^26,27^, we repeated scRNA-seq analysis with a ZC3H20 null *T. brucei* line^27^ to investigate where parasites fail in their development to stumpy forms with respect to transcriptome changes and, potentially, to identify direct or indirect mRNA targets of ZC3H20 itself. We first tested the effect of 10% BHI broth on ZC3H20 null *T. brucei* parasites (ZC3H20 KO)^27^ (Fig. 4a-c). Although the growth rate was reduced in the presence of 10% BHI broth, ZC3H20 KO parasites continued to replicate after WT cells arrested (Fig 4a) and, after 72 hr of culture with 10% BHI, ZC3H20 KO parasites failed to express the PAD1 protein (Fig. 4b). Additionally, we tested the ability of WT and ZC3H20 KO parasites exposed to BHI for 72 hr to differentiate into procyclic cells. Consistent with their inability to generate stumpy forms, after 3 hr of cis-aconitate treatment and incubation at 27 °C, none of ZC3H20 KO parasites expressed EP1 procyclin, in contrast with 84.34% of WT parasites, confirming ZC3H20 KO *T. brucei* fail to differentiate into functional stumpy cells when exposed to BHI broth (Fig. 4c).

**Figure 4.**
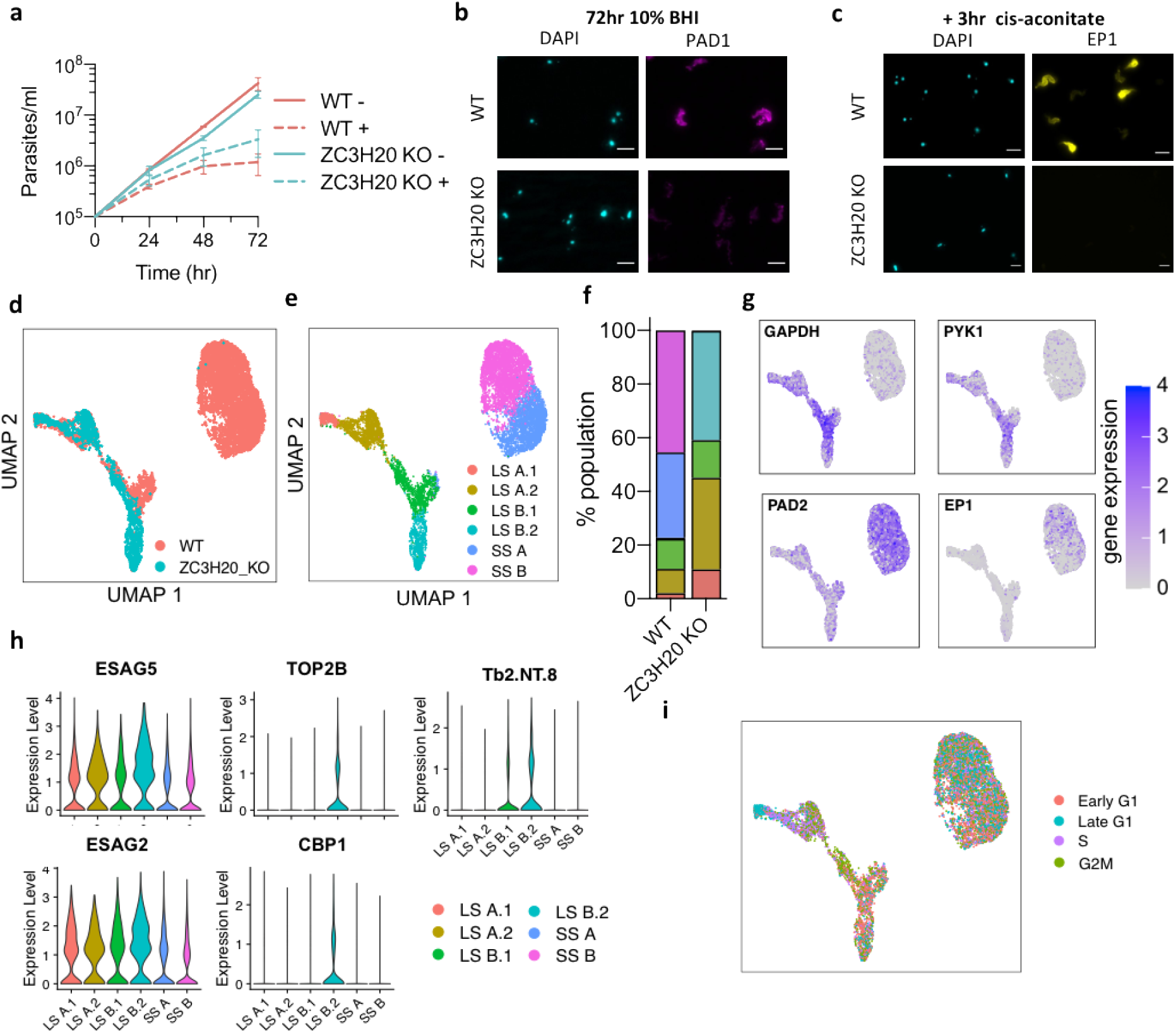
scRNA-seq analysis of differentiation incompetent ZC3H20 KO *T. brucei* parasites. Cumulative growth of WT (red) and ZC3H20 KO (blue) *T. brucei* in culture with (dashed line) and without (solid line) 10% BHI broth. Error bars, SD of three independent replicates. **b**) Staining of WT and ZC3H20 KO parasites with anti-PAD1 antibody after 72 hr incubation with 10% BHI broth. Scale, 5 μm. **c**). EP1 staining of WT and ZC3H20 KO after 3 hr treatment with cis-aconitate to induced differentiation of 72hr BHI + samples into procyclic forms. Scale, 5 μm. UMAP plots of integrated WT cells (red) and ZC3H20 KO cells (blue) coloured by cell type (**d**) and by cluster identification (**e**)**. f**) Proportion of cells in each cluster identified in integrated WT and ZC3H20 KO cells. **g)** UMAP of WT and ZC3H20 KO parasites coloured by transcript count of marker genes as in **1e**. **h**) Violin plots of top slender B.2 marker genes. **i)** UMAP of integrated WT and ZC3H20 KO cells coloured by cell cycle phase.

ZC3H20 KO parasites were next cultured in 10% BHI for 0, 24, 48 or 72 hr and subjected to scRNA-seq, as described for WT samples. After quality control filtering, 2,294 cells (median 1,051 genes per cell) remained and were integrated with the WT cells before dimensional reduction was performed and results were plotted as UMAPs (Fig. 4d, g, h). Clustering the ZC3H20 KO and WT integrated cells resulted in six distinct clusters: stumpy A and stumpy B, and four slender clusters, slender A.1, slender A.2, slender B.1, slender B.2 (Fig. 4d, e). These were identified as slender- and stumpy-like cells by the localised expression of marker genes GAPDH, PYK1, PAD2 and EP1 (Fig. 4g). Whereas 77.3% of WT cells were found in clusters stumpy A or stumpy B, only 0.3% of ZC3H20 KO cells were in either, consistent with near complete ablation of stumpy formation in the mutant parasites (Fig. 4f). The majority of WT slender parasites grouped as members of the slender A.2 and slender B.1 clusters (9.1% and 11.2% of all parasites, respectively), whereas ZC3H20 KO cells were divided between the four slender clusters. Notably, the slender B.2 cohort was comprised almost entirely of ZC3H20 KO parasites (comprising 40.1% of total ZC3H20 KO vs 0.4% of total WT cells). Marker gene analysis between clusters (Fig. S3, Supplementary data 4), identified 94 marker genes upregulated in slender B.2 cells. Of these just 18 genes were uniquely significantly upregulated in cluster slender B.2, the top 5 genes being expression site-associated genes 2 and 5 (ESAG2 and ESAG5), one non-coding RNA gene (Tb2.NT.8), a serine peptidase (Clan SC), Family S10 protein CBP1, and DNA topoisomerase II beta (TOP2B) (fig. 4h). Cell cycle status contributed considerably to clustering analysis of WT and ZC3H20 KO integrated cells, as slender cells clearly group by cell cycle phase (Fig. 4i).

### Trajectory comparison between WT and ZC3H20 KO cells reveals functional separation of downregulation and upregulation of transcripts during differentiation

To compare the transcriptomic changes in ZC3H20 KO and WT *T. brucei* after BHI treatment in detail, we inferred a trajectory from the WT and ZC3H20 KO integrated parasites (Fig. 5a). Doing so identified a branched trajectory: early in pseudotime, WT and ZC3H20 KO parasites were transcriptionally similar and arranged on the same lineage; later, there was a clear branch in their comparative development, ending for WT in stumpy cells and in slender B.2 for ZC3H20 KO cells (Fig. 5a). To understand this change, we first assessed the expression of differentiation-associated genes identified previously in WT parasites (Fig 2d), mapping them across the truncated trajectory branch of the ZC3H20 KO cells (Fig 5b, c). 587 genes of the 2001 identified as differentially expressed during stumpy development in WT cells, significantly changed in expression in ZC3H20 KO parasites across the truncated trajectory (Fig. 5c), although the majority (94.2%) were less highly associated with the ZC3H20 KO trajectory relative to WT (Fig 5d; Supplementary data 4). 75.8% of these genes were part of co-expression modules B-E, which decreased in expression during stumpy development in WT parasites (Fig. 5c). These included genes involved with glycolysis, such as ATP-dependent 6-phosphofructokinase (PFK) and hexokinase 1 (HK1), and the mitotic cell cycle, including cdc2-related kinase 3 (CRK3) and aurora B kinase (AUK1) (fig. 5e). Genes differentially expressed in the ZC3H20 KO trajectory and belonging to expression modules F-H included heat shock 70 kDa protein mitochondrial precursor subunits B and C, and three components of the TCA cycle (succinyl-CoA ligase, mitochondrial malate dehydrogenase and 2-oxoglutarate dehydrogenase E1 component; 2-OGDH E1), but only 2-OGDH E1 increased to a similar level as seen in WT cells (Fig 5e). Hence, ZC3H20 KO cells down regulated transcripts associated with slender cells when exposed to the BHI differentiation stimulus, matching the response of WT cells. However, ZC3H20 KO parasites failed to upregulate transcripts later in development that are required for stumpy formation, and this point of dysregulation coincided with the peak of ZC3H20 expression during normal WT differentiation (Fig. 2g).

**Figure 5.**
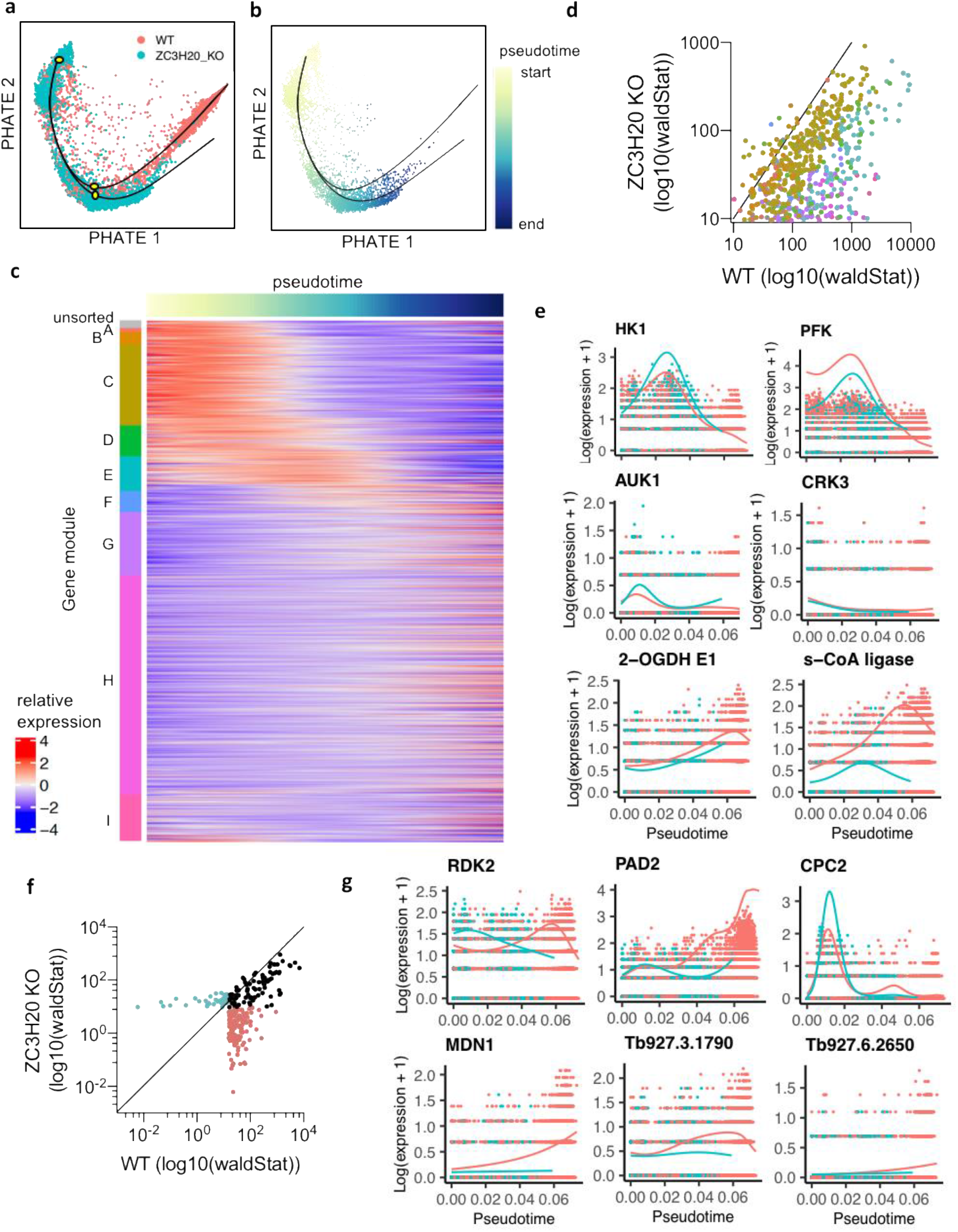
Comparison of differentially expressed genes during differentiation of WT cells and differentiation incompetent ZC3H20 KO cells. **a)** PHATE map of WT (red) and ZC3H20 KO (blue) parasites. Black line indicates branched trajectories. Yellow dots indicate points of analysis for early differentially expressed genes. **b)** PHATE map of ZC3H20 cells only, coloured by pseudotime values assigned for the second lineage of the branch trajectory, black line. **c)** Scatter plot of differentiation-associated genes also found to be differentially expressed in the ZC3H20 KO trajectory. Axes show the association score (log10(wald stat)) for each gene with the WT differentiation trajectory (x-axis) and ZC3H20 KO trajectory (y-axis). Each gene is coloured by is co-expression module identified in **2d**. Black line indicates x=y. **d)** Heatmap of all differentiation associated genes (n = 2001) identified in **2d**. Relative expression across the ZC3H20 KO trajectory is plotted of each gene, grouped by co-expression module. **e)** Expression of example genes across the WT differentiation trajectory (red) and ZC3H20 KO trajectory (blue). X-axis; pseudotime, y-axis; log(expression +1). **f)** Scatter plot of genes identified as early differentially expressed across the branched trajectory (between white points in **a**). Axes show the association score (log10(wald stat)) for each gene with the WT differentiation trajectory (x-axis) and ZC3H20 KO trajectory (y-axis). Genes are coloured by their significant association with the WT (red), ZC3H20 KO (blue), or both trajectories (black). **g)** Expression patterns of example of early differentially expressed genes as in **e**).

To identify regulators of early stumpy development, we looked for genes which changed significantly in abundance from the start of the trajectory to a point just downstream of the ZC3H20 branch (yellow dots, Fig. 5a). 234 genes changed in transcript abundance between these points and were associated with trajectory progression (Fig. 5f, Supplementary data 4). 117 of these genes were associated with both WT and ZC3H20 KO trajectories (p-value <0.05) and include cell cycle associated genes (for example CPC2, Kinetoplastid-specific Protein Phosphatase 1, and structural maintenance of chromosome 4), as expected. Additionally, differentiation associated genes RDK2^28^ and PAD2^35^ were associated with both trajectories but showed different patterns of expression (Fig 5g). 83 genes were differentially expressed only early in the trajectory of WT parasites. These genes include eight relating to ribosome biogenesis (including Midasin, PUF RNA binding protein 10, a putative RNA 3’-terminal phosphate cyclase, and the U3 snoRNA-associated protein UTP11). Other genes include pyruvate dehydrogenase E1 beta subunit, chromatin modifier NLP, TFIIH basal transcription factor complex helicase subunit, Exocyst complex component EXO99, Ras-related proteins RAB7 and RAB2B, and genes encoding hypothetical genes implicated in posttranscriptional regulation of gene expression (Tb927.6.2650, Tb927.11.830 and Tb927.11.7590). 35 genes with early altered expression were associated with the truncated ZC3H20 KO development only. These include G2/M regulator WEE1, kinetoplastid-specific dual specificity phosphatase, Tb927.9.9990, and putative regulator of post-transcriptional gene expression, Tb927.8.3780.

In summary, comparing the differentiation of WT and differentiation incompetent ZC3H20 KO cells through scRNA-seq has allowed the identification of a) the direct and indirect targets of ZC3H20 altered specifically during differentiation, b) the failure point of ZC3H20 KO cells during the temporal profile of differentiation, and c) putative ‘immediate early’ regulators of differentiation.

## Discussion

Although extensively studied, *T. brucei* differentiation from slender to stumpy bloodstream forms has remained difficult to dissect in detail due to the asynchronous nature of this life cycle transition. Here, we used oligopeptide induction of differentiation^23^ in combination with scRNA-seq to deconvolve this process at the transcript level. This approach revealed several details of this process, including: the lack of a discrete intermediate transcriptome; the precise timing of cell cycle exit, immediately prior to late G1; the transient expression of several genes not identified by bulk-analysis; and the expression timing of known and putative differentiation factors during the developmental processes. Using scRNA-seq to study ZC3H20 KO parasites, we were also able to validate the essentiality of ZC3H20 for differentiation and position its action specifically at the major slender to stumpy transition point where the transcripts of this gene peak in abundance. Additionally, we were able to provide detailed gene expression patterns of both known and novel cell cycle regulated genes during the slender cell cycle.

Clustering WT *T. brucei* into groups of transcriptionally similar cells clearly identified two primary groups of slender- and stumpy parasites, each of which could be further classified into two sub-slender and sub-stumpy clusters (Fig 1). The transcript differences between clusters were mainly due to the cell cycle phase of slender cells and the stage of progression towards stumpy development (Fig 1 and 2) with gene expression changes highlighting a progressive transition to stumpy forms with relatively few genes transiently changing in abundance during development (n = 96, FC >2). Thus, although discrimination of parasites between the extremes of the slender and slender morphotypes is possible microscopically, scRNA-seq analysis does not provide evidence for an intermediate form defined in molecular terms. Rather, short stumpy cells appear to emerge directly from the G1 phase of replicative slender cells (see below).

Previous bulk transcriptomics identified marker genes of the *T. brucei* cell cycle phases (early and late G1, S phase and G2/mitosis)^69^, allowing us to define the most likely cell cycle position of each cell (Fig 3). Phase identification of the cycling parasites showed that late G1 stage cells are positioned at the start of the differentiation trajectory, consistent with a cell cycle receptive window^74^, followed by S and G2/M phase cells. Early G1 cells were enriched closer to stumpy clusters, though less clearly grouped, and stumpy clusters showed no clear cell cycle phase (Fig. 3, 4i, and 4j). Hence, the major switch in transcriptome from slender to stumpy occurs during G1 and, specifically, before cells enter late G1. Cells at other stages may be committed to differentiation but not yet arrested, as predicted by modelling^75^. The inability to assign stumpy cells to any one proliferative cell cycle phase indicates these cells persist in a distinct G0, as opposed to simply pausing in G1. The signalled progression of *T. brucei* into G0 as stumpy forms provides a valuable, evolutionary divergent and tractable model for studying the conservation of quiescence signalling pathways, which are critical in many eukaryotic developmental processes^76–80^.

We were additionally able to investigate the changes in transcript abundance during the proliferative slender cell cycle. Pseudotime analysis allowed us to profile the dynamic patterns of 1,897 genes found to be differentially expressed (Fig. 3), 332 of which had been previously identified in bulk RNA-seq analysis of synchronised procyclic *T. brucei*^69^. For example, we find CYC4, CYC6 and CYC8 peak at distinct points: CYC4 is a CYC2-like protein predicted to act in G1^81^, where the transcript levels increase in our analysis; CYC6^62,82,83^ is known to regulate nuclear division and peaks post S-phase; and predicted, but untested, mitotic cyclin CYC8 peaks very precisely in late G1/S phase cells. Having not undergone either selectional or chemical synchronisation procedures^69,84,85^, this scRNA-seq derived cell cycle atlas provides a relatively unperturbed picture of cell cycle regulated events in greater detail than previously available, and suggests candidates for functional analysis. Distinctions between developmentally competent (pleomorphic) slender forms and adapted monomorphic forms used in previous studies may also be identified.

Trajectory inference and differential expression analysis of the slender to stumpy transition revealed the relative order of events during differentiation of this asynchronous population. Initially, there was higher abundance of transcripts linked to proliferation (Fig. 1 and 2), with cells completing the later stages of the cell cycle during the early stages of the differentiation trajectory, consistent with phase scoring analysis (Fig. 2 and 3). Thereafter, metabolic changes and activation of the mitochondrion occurred as expected^12,38^. Finally expression of several kDNA encoded genes (cytochrome oxidase subunits I-III and cytochrome B) and procyclin surface protein encoding genes, EP1, EP2 and GPEET was observed, reflecting preparation for differentiation to procyclic forms^12,15,17^. These changes correlated well with bulk mRNA analysis of *in vivo* parasites^12^, validating the use of BHI as an *in vitro* model of stumpy development. Transient expression patterns of several genes, not discernible in bulk RNA-seq and proteomic studies, were also observed. These included several hypothetical genes, which may prove to be negative or positive regulators of differentiation. For instance, Tb927.8.7820 peaked precisely at the slender to stumpy transition point, is able to decrease mRNA stability^68^ and is negatively targeted by RBP10^86^, which in our analysis decreased during differentiation consistent with its reported function in maintaining the bloodstream cell state^39,40,86^. Other transiently regulated transcripts were associated with cell cycle or mitochondrial control, including CYC6, required for nuclear division^62,82,83^, CIF1, a master regulator of cytokinesis^43^ and MtHSP70B and MtHSP70C, which locate to the mitochondrial outer membrane^87^ and may facilitating protein folding and targeting in the developing mitochondrion, matching the function of the human homolog, HSPA9^88^.

Transiently upregulated genes also included the known differentiation regulator ZC3H20^26,27,46^, confirming that we were able to identify developmental regulators via their gene expression patterns. We postulated that scRNA-seq analysis may enable us to map cells with diverse differentiation phenotypes onto our trajectory of WT differentiation, to assess the point at which genetically perturbed parasites fail to develop. We therefore exposed ZC3H20 KO parasites^27^ to oligopeptides and confirmed that they remained proliferative (Fig. 4i, j) and failed to develop to stumpy forms (Fig. 4 d-h). Trajectory inference revealed that ZC3H20 KO cells downregulate transcripts also downregulated in oligopeptide stimulated WT parasites, including several glycolysis factors, cell cycle regulating genes, and posttranscriptional regulators of gene expression (Fig. 5). This downregulation may contribute to the reduced growth of ZC3H20 KO parasites when exposed to BHI. Interestingly, however, there was a clear distinction between this downregulation of slender transcripts and the increase of stumpy-associated transcripts, which ZC3H20 KO parasites failed to upregulate to WT levels, including 12/19 ZC3H20 regulated mRNAs^26^. This suggests that ZC3H20 KO parasites perceive the differentiation signal and undergo early steps of differentiation, but do not commit to cell cycle exit and further development to stumpy forms. Comparing the divergence of WT and ZC3H20 KO cells in the trajectory of differentiation identified putative ‘immediate early’ regulators of commitment (Fig. F), including 28 hypothetical genes, four of which are known to regulate mRNA stability^68^. Further experimental work will be required to test the involvement of these genes in differentiation.

In summary, our data demonstrate that transcript level changes in parasites could be used to compile maps of both the cell cycle and the asynchronous slender to stumpy differentiation process. These can be mined to identify regulatory genes of individual events that make up each process. We further characterised mutant parasites by the same approach, positioning the site of action of one regulator (ZC3H20) in the developmental time course. If iterated for different genes, this method can be exploited to derive hierarchies of gene action during differentiation in this and other life cycle stages, species and development processes.

## Materials and methods

### *Trypanosoma brucei* cell lines and culture

Trypanosoma brucei EATRO 1125 AnTat1.1 90:13 parasites^89^ were used as pleomorphic wild-type (WT) in all experiments. The ZC3H20 KO null parasites were previously generated in the same cell line transfected with plasmid pJ1399 (gifted by Dr. Jack Sunter), containing T7 polymerase and CRISPR/cas9, by replacement of both alleles of Tb927.7.2660 with blasticidin S deaminase^27^. All parasites were grown free from selective drugs in HMI-9 medium^90^ (Life technologies), supplemented with 10% foetal calf serum at 37°C, 5% CO2. For induction of differentiation, parasites were maintained below ~7×10^5^ cells per ml for up to 5 days prior to addition of brain heart infusion (BHI) broth (Sigma Aldrich).

### Single cell RNA-sequencing

For each scRNA-seq sample, four staggered cultures were set up over four days all maintained below ~8×10^5^ cells per ml during the experiment by dilution. One culture was maintained free from BHI, and the remaining had 10% BHI added 24, 48 or 72 hr prior to sample preparation. Equal numbers of parasite from each culture were then combined to generate one pooled sample. 1.5 ml of the pooled culture was centrifuged, and the pelleted cells washed twice with ice-cold 1 ml 1X PBS supplemented with 1% D-glucose (PSG) and 0.04% Bovine Serum Albumin (BSA). Cells were then resuspended in ~ 500 ul PSG + 0.04% BSA, filtered with 40 μm Flowmi™ Tip Strainer (Merck) and adjusted to 1,000 cells/μl. In all steps, cells were centrifuged at 400 x g for 10 minutes. 15,000 cells (15 μl) from the mixed sample were loaded into the Chromium Controller (10x Genomics) to capture individual cells with unique barcoded beads. Libraries were prepared using the Chromium Single Cell 3ʹ GEM, Library & Gel Bead Kit v3 (10x Genomics). Sequencing was performed with the NextSeq™ 500 platform (Illumia) to a depth of ~50,000 reads per cell. Library preparation and sequencing was performed by Glasgow Polyomics. For the first WT replicate experiment, *T. brucei* parasites were mixed 1:1 with *Leishmania mexicana* prepared by the same method (data unpublished), so the heterogenous doublet rate of 8.04% could be calculated.

### Read mapping and transcript counting

The reference genome was complied with Cell Ranger, to combined the TREU927 *T. brucei* nuclear reference genome^91^ and *T. brucei* EATRO 1125 maxicircle kDNA sequence^67^. 3’ UTR annotations were extended to increase the proportion of reads correctly assigned to annotated transcripts. 2500 bp immediately downstream of the stop code was assigned as the 3’UTR of each protein coding gene, unless the existing 3’UTR was longer than 2500 UTR in which case the full length was preserved. If the new 3’UTR was overlapped with other genome features (coding and non-coding) the UTR was truncated to remove the overlap. Reads were mapped and unique reads aligned to each annotated gene were counts and assigned to a cell barcode with the Cell Ranger count function (Supplementary data 1). Cell Ranger v3.0.2 (http://software.10xgenomics.com/single-cell/overview/welcome) was used with all default settings.

### Data processing and integration

Count data for individual samples (WT 1, WT 2 and ZC3H20 KO) was processed separately prior to integration using the Seurat v3^92^ and Scran v1.14.5^93^ packages with R v3.6.1. The percentage of transcripts encoded on the maxi circle kDNA was calculated per cell, as cells with excess proportion of mitochondrial transcripts are likely to be poor quality^94^. The percentage of transcripts per cell encoding ribosomal RNA was also calculated, as high levels of rRNA indicate poor capture of polyadenylated transcripts. Low quality cells were removed by filtering for low total RNA (<1,000), low unique transcripts (< 250), high proportion of kDNA (> 2%) and high proportion of rRNA (>8%). Likely doublets were removed by filtering for high total RNA (> 4,000) and high total unique transcripts counts (> 2,500). After filtering rRNA transcripts were removed from each cell’s transcriptome. For sample metrics, see Supplementary data 1.

Each filtered sample was log normalised individually using the quick cluster method from Scran^95^. To increase the robustness of variable genes selected for principle component (PC) analysis, we used two selection methods^96^; Scran, which uses log normalised transcript counts, and Seurat^92^, which uses raw transcript counts. We identified 3,000 genes with each method, selected those identified by both and removed VSG encoding genes^97^ to avoid clustering based on VSG expression. This left 2,029, 1,643 and 2,132 for WT 1, WT 2 and ZC3H20 KO samples, respectively (Supplementary data 1).

For integration of WT replicate samples, the Seurat v3 package was used^92^. Common variable features and integration anchors were identified, data for all genes integrated and scaled before the PCs were calculated using the common variable features. The first 8 PC dimensions each contributed > 0.1% of additional variance and were used to select anchors and integrate data. The effect of total RNA per cell was regressed when scaling data. The ZC3H20 KO cells were subsequently integrated with the previously integrated WT data using the steps described above, however STACAS v1.01.1 was used to identify integration anchors as the package is specialist for samples which don’t fully overlap^98^.

### Cluster analysis and mark gene identification

For clustering and marker gene analysis the Seurat v3 package was used^92^. Cells were plotted as dimensionality reduced UMAPs^37^ and nearest neighbours were identified using 8 dimensions. A range of clustering resolutions were trialled, with 0.4 resulting in the highest resolution clustering with significant mark genes identified for every cluster. Marker genes were identified for each cluster using MAST^99^. Only genes expressed in > 25% of the cells in the cluster, with a logFC of > 0.25 and adjusted p-value < 0.05 were considered marker genes. Gene ontology (GO) terms concerning biological processes were identified via the TriTrypDB^100^ website (p <0.05) and redundant terms removed with REVIGO^101^ (allowed similarity = 0.5) and manually.

### Trajectory inference and pseudotime analysis

For trajectory inference cells were plotted using PHATE maps^60^ (using the same common viable genes as for PCA and 8 dimensions) and trajectories were identified using slingshot^102^, with the slender A cluster defined as the starting point. For cell cycle analysis, a circular trajectory was fitted as a principle curve^103^. To identify genes with expression patterns associated with progression of the trajectory, generalise additive models were fit using the tradeSeq package v1.3.18^104^ with default parameters. The number of knots was tested to find 6 knots provide sufficient detail for the highest number of genes without overfitting. Differential expression analysis was performed with the tradeSeq associationTest function using default parameters, and significant genes (p-value <0.05, FC >2) were clustered using tradeSeq clusterExpressionPattern over 100 points on the trajectory. Gene clusters were merged into co-expressed modules using default setting except the merging cut-off was set to 0.95 to refine the number of modules from 58 to 9. For comparison with bulk RNA-seq analysis, the fold-change of stumpy vs slender expression for all genes was taken from data published by Silvester *et al.*^12^. All genes not found to be significant (p-value > 0.05) in bulk analysis were given a fold-change value of 0 for comparison with scRNA-seq data.

### Immunofluorescence

Parasites were fixed in 1% paraformaldehyde for 10 minutes at room temperature (RT). Parasites were washed in 1X PBS and adhered to slide spread with Poly-L-lysine before being permeabilised with 0.1% Igepal in 1X PBS for 3 minutes. Cells were then blocked with 2% BSA in 1X PBS for 45 minutes at RT, stained with primary antibody (anti-PAD1^35^ 1:1,000, EP1 procyclin [Cedar labs] 1:300) diluted in 0.2% BSA for 1 hr at RT. Three washes with 1X PBS were performed before incubating with secondary Alexa Fluor 488 (ThermoFisher Scientific) in 0.2% BSA for 1 hr at RT. Cells were washed a further three times before mounting with Fluoromount G with DAPI (Cambridge Bioscience, Southern Biotech). Imaging was performed with an Axioscope 2 fluorescence microscope (Zeiss) and a Zeiss Plan Apochromat 63x/1.40 oil objective.

## Data availability

Data can be sources via Supplementary Data Tables and the European Nucleotide Archive with accession number PRJEB41744. Wild-type scRNA-seq data can be explored using the interactive cell atlas (http://cellatlas.mvls.gla.ac.uk/TbruceiBSF/).

## Code availability

Code used to perform analysis described can be accessed at GitHub (https://github.com/emma23ed/Tbrucei_scRNA-seq.git).

## Extended data

Supplementary Data 1. scRNA-seq sample metrics and variable genes.

Supplementary Data 2. Analysis of wild-type differentiating *T. brucei*

Supplementary Data 3. Cell cycle analysis of slender form *T. brucei*

Supplementary Data 4. Analysis ZC3H20 KO *T. brucei*

## Acknowledgements

We thank J. Galbraith and P. Herzyk (Glasgow Polyomics, University of Glasgow) for their guidance, library preparation and sequencing. We also thank F. Rojas and M. Cayla (University of Edinburgh) for their guidance and provision of cell lines. This work was supported by the Wellcome Trust (218648/Z/19/Z to E.M.B., 104111/Z/14/ZR to T.D.O. and 103740/Z14/Z to K.R.M.), Wellcome Trust Institutional Strategic Support Fund (ISSF3) awards held at the University of Glasgow (204820/Z/16/Z awarded to E.M.B. and R.M.), and the BBSRC-FAPESP (BB/N016165/1 to R.M).

## Author Contributions

Methodology: E.M.B., R.M., T.D.O, and K.R.M. Data collection: E.M.B. Bioinformatic data analysis: E.M.B and T.D.O. Single cell atlas was created by T.D.O. All authors participated in discussions related to this work. All authors wrote, reviewed and approved the manuscript.

---

**Fig S1.**
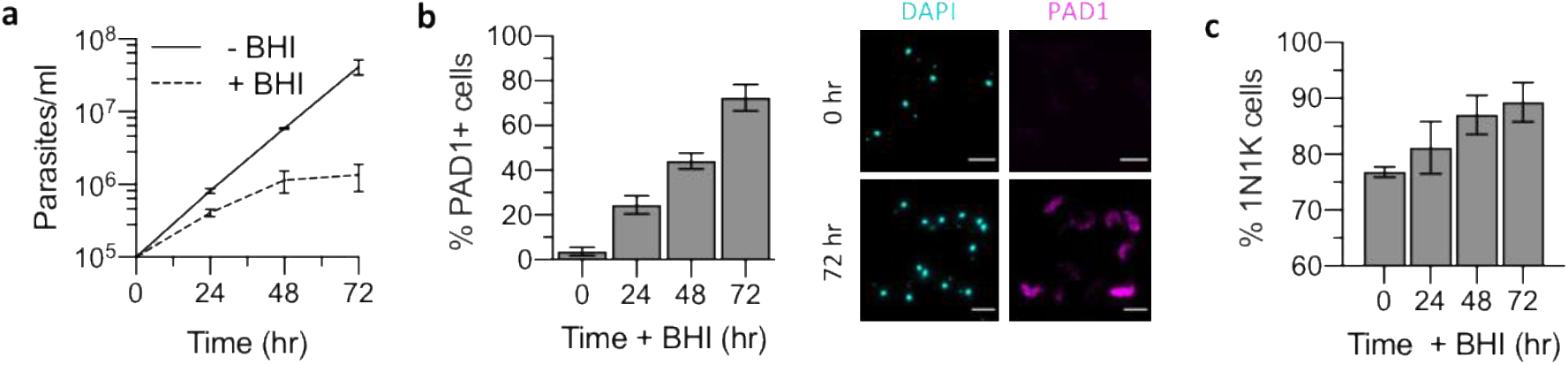
Brain heat infusion (BHI) broth induces slender to stumpy differentiation *in vitro.* **a**) Cumulative growth of pleomorphic *T. brucei* in culture with (dashed line) and without (solid line) 10% BHI broth. Error bars, SD of three independent replicates. **b**) Left: Percentage of cells expressing stumpy marker protein PAD1 after culturing with 10% BHI. Error bars, SD of triplicate samples. Right: Staining of parasites with anti-PAD1 antibody, after 72 hr incubation with 10% BHI broth. Scale, 5 μm. **c**) Percentage of 1N1K *T. brucei* after culturing with 10% BHI broth. Error bars, SD of three independent experiments.

**Fig S2.**
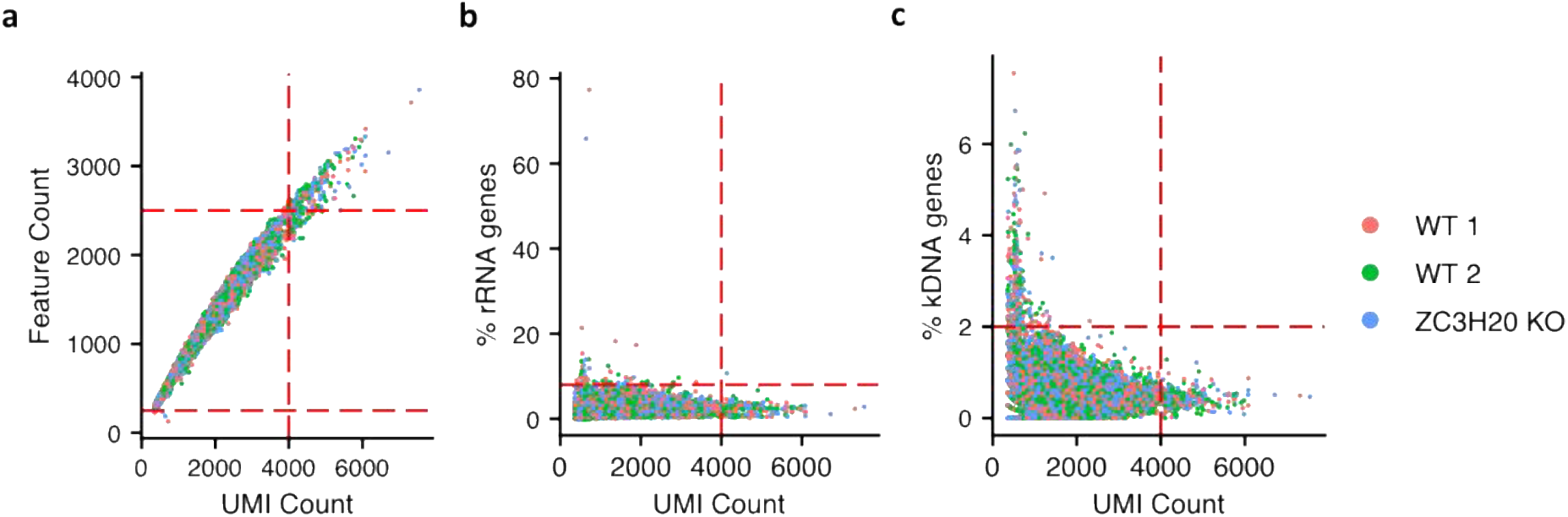
Quality control and filtering of single transcriptomes. Scatter plots for WT replicate 1(red) and 2 (green), ZC3H20 KO (blue) experiments; each data point is one transcriptome. Plots show the relationship between the number of unique molecular identifiers detected per cell and (**a**) number of features (genes), (**b**) percentage of features encoding ribosomal RNA (rRNA) and (**c**) percentage of features encoding on the kDNA maxi circle genome. Red dashed lines indicate thresholds used to filter cells per experiment; Features count > 250, < 2500, UMI count < 4000, % rRNA genes < 8, % kDNA genes < 2.

**Fig S3.**
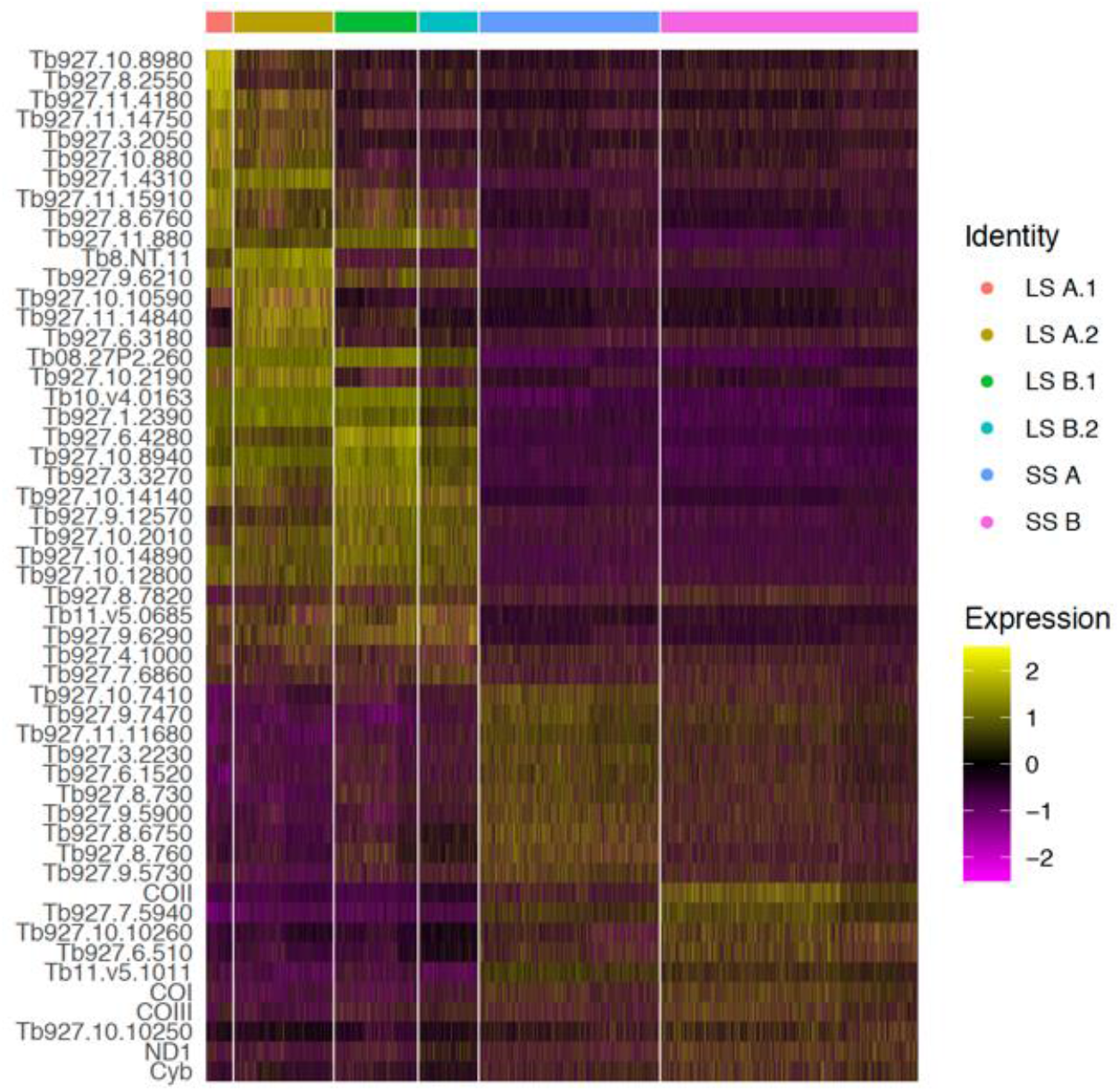
WT and ZC3H20 KO cluster maker genes. Heatmap showing relative expression of the top 10 unique maker genes of each cluster identified in **4d**. Each row is one gene coloured by relative expression. The gene ID is given for each marker. Each column is one cell grouped according to cluster identity.

